# Soil stabilisatizion for DNA metabarcoding of plants and fungi. Implications for sampling at remote locations or via third-parties

**DOI:** 10.1101/2020.09.07.280016

**Authors:** Lina A Clasen, Andrew P Detheridge, John Scullion, Gareth W Griffith

**Affiliations:** Institute of Biological, Environmental and Rural Sciences, Aberystwyth University, Adeilad Cledwyn, Penglais, Aberystwyth, Ceredigion SY23 3DD, WALES

**Keywords:** Freeze-drying, chitinolytic fungi, Freeze-thaw, sample preservation

## Abstract

Storage of soil samples prior to metagenomic analysis presents a problem. If field sites are remote or if samples are collected by third parties, transport to analytical laboratories may take several days or even weeks. The bulk of such samples and requirement for later homogenisation precludes the convenient use of a stabilisation buffer, so samples are usually cooled or frozen during transit. There has been limited testing of the most appropriate storage methods for later study of soil organisms by eDNA approaches. Here we tested a range of storage methods on two contrasting soils, comparing these methods to the control of freezing at −80°C followed by freeze-drying. To our knowledge this is the first study to examine the effect of storage conditions on eukaryote DNA in soil, including both viable organisms (fungi) and DNA contained within dying/dead tissues (plants). For fungi, the best storage regimes (closest to the control) were storage a 4°C (for up to 14 d) or active air-drying at room temperature. The worst treatments involved initial freezing followed by thawing which led to significant later spoilage. The key spoilage organisms were identified as *Metarhizium carneum* and *Mortierella* spp., with a general increase in saprotrophic fungi and reduced abundances of mycorrhizal/biotrophic fungi. Plant data showed a similar pattern but with greater variability in community structure especially in the freeze-thaw treatments, probably due to stochastic variation in substrates for fungal decomposition, algal proliferation and some seed germination. In the absence of freeze drying facilities, samples should be shipped refrigerated but not frozen if there is any risk of thawing.

## Introduction

The use of eDNA metabarcoding (amplicon sequencing) has transformed our knowledge of the structure and composition of soil biological communities (Geml et al., 2014; Williams, 2020), with more recent metagenomic studies enhancing our understanding of the metabolic processes mediated by these organisms (Keepers et al., 2019; Ogwu et al., 2019). However, the methods used to sample soils (Epp et al., 2012; Lindahl et al., 2013; Taberlet et al., 2012), store and extract total soil DNA/RNA (Kennedy et al., 2014; Soliman et al., 2017) can exert a strong influence of the data obtained, as can the barcoding loci/primers and sequencing platforms used. Thus there is a need for a standard operating procedure (SOP) before metabarcoding analyses conducted in different laboratories can be compared (Lindahl et al., 2013; Orgiazzi et al., 2015).

As the use of eDNA metabarcoding has extended to the study of soils in more remote locations (Detheridge et al., 2020; Tedersoo et al., 2014) and to more applied deployment in nature conservation site monitoring by statutory organisations (Detheridge et al., 2018; Geml et al., 2014; Latch, 2020; Valentin et al., 2020), the issue of soil storage between sampling and subsequent analysis has become an important consideration. Such concerns have interested soil scientists for many decades but usually in relation to the metabolic status of soil organisms, assessed via community level physiological profiles (CLPP), soil respiration etc. (Lee et al., 2007). However, for metabarcoding / metagenomic studies, the preservation of DNA/RNA unchanged from the natural state presents its own distinctive challenges.

There are established international guidelines recommending refrigerated storage or freezing (OECD, 2000) when chemical or microbiological analyses cannot be undertaken on fresh soil. However, the preservation of soils by air-drying dates back to the origins of soil science and remains the simplest method for long-term stabilisation. Examination of air-dried soil archive samples dating back over a century by Clark et al. (2008) found that whilst long-term storage of air-dried soil reduced the amount of DNA present, differences in bacterial populations according to soil plot treatment were still detectable.

Current standards have been established through the International Organization for Standardization (ISO), for example ISO 11063 (2012) (“Soil quality — Method to directly extract DNA from soil samples”) and ISO-10381-6 (2002) (“Soil quality—Sampling—Part 6: Guidance on the collection, handling and storage of soil for the assessment of aerobic microbial processes in the laboratory”). However, these are focused predominantly on bacteria (Petric et al., 2011), which appear to respond in storage rather differently from fungi and other groups of biota (Terrat et al., 2015). However, it is unclear whether this is due to the greater tolerance of bacterial cells to disruption by freezing etc. or the lower taxonomic resolution of the standard 16S metabarcoding procedures.

Martí et al. (2012) examined the effects of different soil storage and found that the stability of bacterial DNA (assessed via DGGE) varied according to soil type. Similarly Lauber et al. (2010), used DNA metabarcoding of bacterial populations to examine the effects of storage of soil or faeces at a range of temperatures (−80°C, −20°C, 4°C, 20°C) for 3 or 14 days. Even after 14 d at 20°C (in sealed container), they found only small changes in bacterial communities. However, Rubin et al. (2013), also using DNA metabarcoding to assess changes in soil bacterial populations, found that there was a progressive loss of diversity associated with storage under warmer conditions.

Freeze drying (with subsequent frozen storage) is widely considered by many to be the best available option for the stabilisation of soils prior to nuclei acid extraction (Castaño et al., 2016; Straube and Juen, 2013; Weißbecker et al., 2017). Initial freezing inactivates biological processes and later removal of water via sublimation at low pressure, stabilises the soil in an inactive dry state. However, Bainard et al. (2010) found a reduction in arbuscular mycorrhizal fungal DNA in roots following long-term storage of freeze-dried roots at ambient temperature, so frozen storage following initial processing is important here too. There are significant additional advantages to freeze-drying. Freshly-collected soils can be frozen immediately without prior processing and after freeze-drying stored frozen in a stable state, convenient (unlike directly frozen soil) for later homogenisation/grinding. The ease with which freeze-dried soils can be finely ground, allows for efficient homogenisation of larger samples. The latter point is important, since it is crucial that subsampling for DNA extraction (commonly 200 mg from 500g samples) is representative of the whole (Lindahl et al., 2013). Repeated DNA extractions from a fully homogenised soil would therefore result in the same community structure derived from metabarcoding. However, availability of suitably large freeze-drying capacity can be an important limiting factor at many institutions.

Where it is not possible to freeze samples within a few hours of collection, the question remains as to what pre-treatment is best to preserve the nucleic acids of the soil communities during shipping from field sites (often sampled by third parties) to analytical labs. Here we compare the effect of immediate freezing to a range of different soil DNA stabilisation methods, using equipment available outside laboratories (freezers, fridges, fans and ovens). The resulting effects of these soil storage methods are examined using eDNA metabarcoding profiles for plants and fungi, hypothesising that inferior storage conditions would lead to: a general loss of diversity in both plants and fungi, due to DNA degradation; proliferation of a subpopulation of faster-growing fungi, well-suited to growth in particular storage conditions, which would be associated with greater levels of DNA degradation.

## Methods

Soil was collected from an upland (grazed) grassland immediately adjacent to the Brignant longterm grazing experimental field site (lat/long: 52.3648°N, 3.8214°W; 367m asl.). Brignant soil is an acidic (pH 5) Manod Series (loam over shale Palaeozoic slate, mudstone and siltstone; well-drained fine loamy or fine silty soils over rock; (Hallett et al., 2017)) with an organic carbon content of 7.3%. Turf was removed to a depth of 3-5 cm from a single 0.25 m^2^ area and approximately 10 kg of soil collected from the remaining 10 cm depth of topsoil avoiding large stones. A contrasting less acidic and lower organic matter, alluvial soil type (Conway series 0811b [Fluvic Eutric Gleysols] groundwater gley (silt to silty-clay loam. Depth to gley layer varies @ 10 - 20 cm.)(Hallett et al., 2017)) pH 6.1, organic carbon 3.6%, was also collected (15 kg) from the edge of an arable field at Gogerddan (52.4364°N, 4.0313°W; 15 m a.s.l.), with removal of vegetation, as above, and processed in the same way. The soils were transported within 2 h to the laboratory and sieved (3 mm) to remove further stones and to enable thorough homogenisation of the soil/roots. Samples of 200 g were weighed into Ziploc bags (40 in total) and divided into treatments with 4 replicates per treatment. Bags for the control treatment (Treatment 1) were immediately frozen at −80°C.

The soil treatments are shown in Table 1. Treatments T2, T3 and T4 were designed to test the effect of initial freezing followed by thawing during shipping either refrigerated for 14 d (T2) or 5 d (T3), or stored at ambient temperature (23°C, T4). Treatments T5, T6 and T7 were all dried for 5 d after an initial “shipping” period (storage for 3d at 4°C). T5 was dried more rapidly by blowing ambient air into open bags (hairdrier but no heat; as shown in SuppFig 1), whereas T6 and T7 were passively dried in open bags at room temperature and 37°C respectively. Treatments T8, T9 and T10 involved storage in closed bags either at 4°C for 14 d (T8) or at ambient temperature (T9 for 14 d and T10 for 5 d). Five of these 10 treatments (T1, T3, T6, T8 and T9) were also applied to the alluvial soil to see if a contrasting soil responded to storage in a similar manner.

**Table 1.**
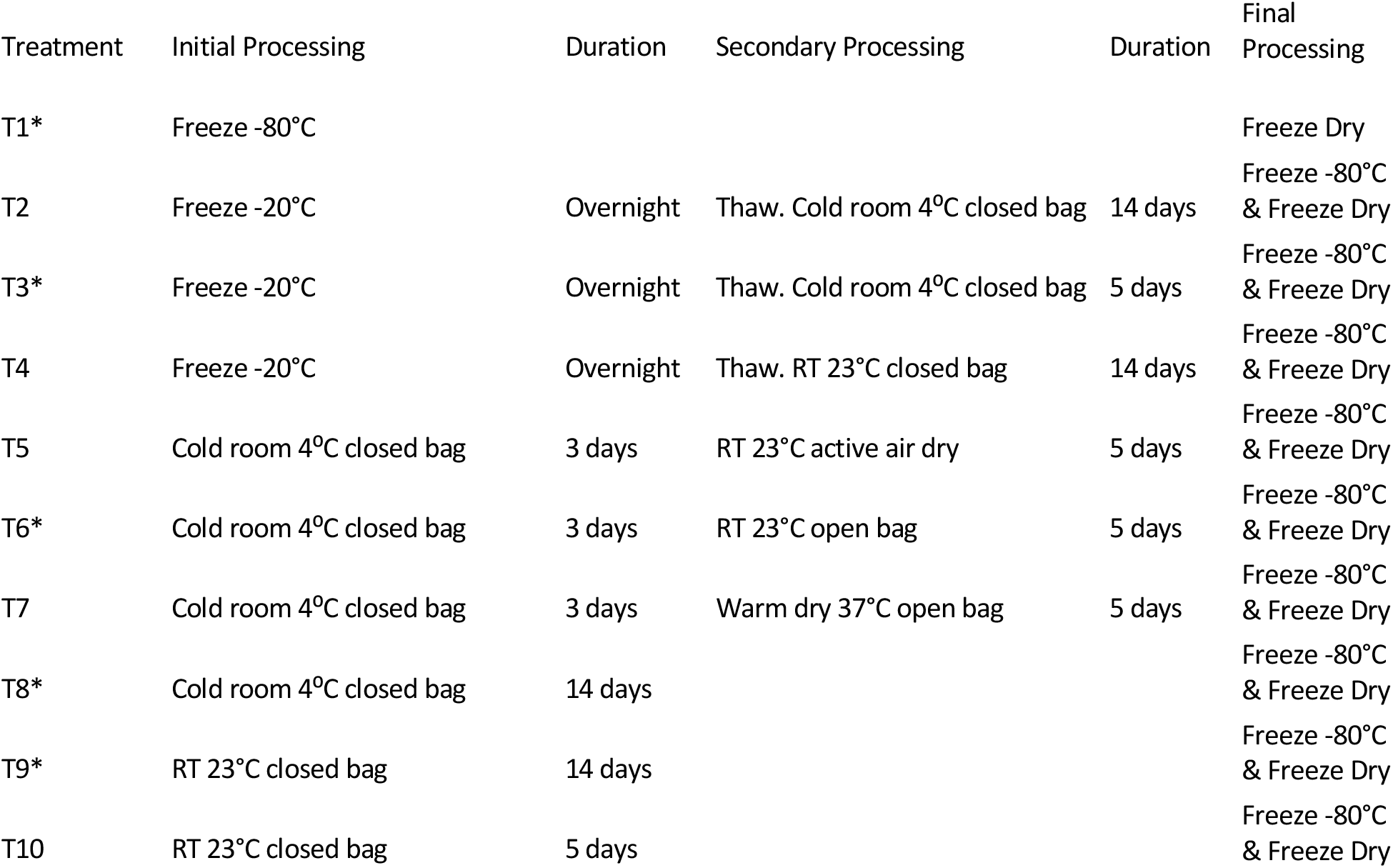
Storage treatments tested in this study using soil adjacent to the Brignant longterm experiment. An * indicates treatments also tested with soil from the Gogerddan alluvial plain.

| Treatment | Initial Processing | Duration | Secondary Processing | Duration | Final Processing |
| --- | --- | --- | --- | --- | --- |
| T1* | Freeze -80°C |  |  |  | Freeze Dry |
| T2 | Freeze -20°C | Overnight | Thaw. Cold room 4°C closed bag | 14 days | Freeze -80°C & Freeze Dry |
| T3* | Freeze -20°C | Overnight | Thaw. Cold room 4°C closed bag | 5 days | Freeze -80°C & Freeze Dry |
| T4 | Freeze -20°C | Overnight | Thaw. RT 23°C closed bag | 14 days | Freeze -80°C & Freeze Dry |
| T5 | Cold room 4°C closed bag | 3 days | RT 23°C active air dry | 5 days | Freeze -80°C & Freeze Dry |
| T6* | Cold room 4°C closed bag | 3 days | RT 23°C open bag | 5 days | Freeze -80°C & Freeze Dry |
| T7 | Cold room 4°C closed bag | 3 days | Warm dry 37°C open bag | 5 days | Freeze -80°C & Freeze Dry |
| T8* | Cold room 4°C closed bag | 14 days |  |  | Freeze -80°C & Freeze Dry |
| T9* | RT 23°C closed bag | 14 days |  |  | Freeze -80°C & Freeze Dry |
| T10 | RT 23°C closed bag | 5 days |  |  | Freeze -80°C & Freeze Dry |

**SuppData 1**. Photo of the hairdrier apparatus used for active air-drying of soil (treatment T5)

After the storage treatments were completed, all bags were frozen at −80°C. Samples were then freeze-dried before sieving at 0.5 mm and thoroughly homogenised according to our standard lab procedure (Detheridge et al., 2016). DNA was extracted from 200mg of freeze dried soil using the Power Soil DNA extraction kit (Qiagen), as described by Detheridge et al. (2016). The ITS2 region of plants and fungi were amplified using a mix of primers. For fungi, the forward primers were those used by Tedersoo et al. (2014), with an equimolar mix of 6 primers (SuppData 2). To this mix the plant primer Chen S2F (ATGCGATACTTGGTGTGAAT) (Chen et al., 2010) was added in a ratio of 3 fungal mix to 1 plant primer. This ratio was chosen to ensure that the majority of sequences returned were fungal as this is the prime aim of the analysis and fungal communities are generally more complex than plant communities. The reverse primer was the universal ITS4 primer. The forward primers were linked at the 5’ end to the Ion Torrent B adapter sequence (CCTCTCTATGGGCAGTCGGTGAT). The ITS4 primer was linked at the 5’ end to the Ion Torrent A-adapter sequence (CCATCTCATCCCTGCGTGTCTCCGAC), the TCAG key and an IonXpress Barcode.

**SuppData 2**. Fungal forward primer sequences with target group to amplify all fungal groups, and also Stramenopiles (Ooomyces) devised by Tedersoo et al. (2014)

PCR was carried out using PCR Biosystems Ultra polymerase mix (PCR Biosystems Ltd, London UK). Each reaction contained 250 nM of the forward primer mix and 250 nM of reverse primer. Amplification conditions were initial denaturing 15 min at 95°C, followed by 30 cycles at 95°C for 30 s, 55°C annealing for 30 s, 72°C extension for 30 s, and a final extension of 5 min at 72°C.

After PCR, samples were processed, sequenced and sequence data processed as detailed in Detheridge et al. (2016; 2018; 2020). Fungal sequences were identified using a database build from v8.0 of UNITE (Abarenkov et al., 2019) and plants sequences using a database built as detailed in Detheridge et al. (2020). Sequence data have been submitted to the European Nucleotide Archive with reference number XXXXXXX. Data were expressed as relative abundance of each of the species detected (separately for plants and fungi).

Principal coordinate ordination (PCO) visualised differences in community structure using square root transformed abundances and a Bray-Curtis distance matrix; these analyses were undertaken in R (R_Core_Team, 2013). PERMANOVA determined whether there were significant differences in fungal and plant communities between treatments and the pairwise test used to determine which treatments differed and their degree of separation. An analysis of similarity (SIMPER) was used to determine which OTUs varied between treatments. These analyses were conducted in PRIMER-PERMANOVA + v6. ANOVA was used to determine the significance of treatment effects on relative abundance and these were carried (in R), after any appropriate transformations to meet requirements of analyses.

## Results

After quality checking there were a total of 2 772 707 ITS2 sequences with a maximum of 94 109 sequences per sample and a minimum of 52 931 (Mean 69 318). After rarefying to the lowest number of sequences per sample, dropping singleton sequences and trimming 5.8S and 28S regions, clustering resulted in 848 plant and fungal OTUs for the upland (Brignant) soil and 769 for the alluvial (Gogerddan) soil.

### Upland (Brignant) soil

Principal coordinate ordinations of the fungal community data (Fig 1 A), show the clear outlier relative to T1 (control) is T4 (Freeze-thaw, left at 23°C for 14 d), with T2/T3 (Freeze-thaw, left at 4°C for 5 d/14 d respectively) and T7 (storage at 4°C followed by passive warm air-drying at 37°C) being the next most divergent treatments. PERMANOVA analyses showed a significant effect of storage treatment for the fungal community data (Pseudo F = 5.1728 P= 0.001). Pairwise Permanova comparisons of the fungal populations in each treatment with the control (Fig. 2A) confirm that treatments T2, T3, T4, and T7 had a significant effect on the fungal populations present at the end of the storage period. However, for the other pre-treatments there were no significant differences relative to control.

**Fig. 1.**
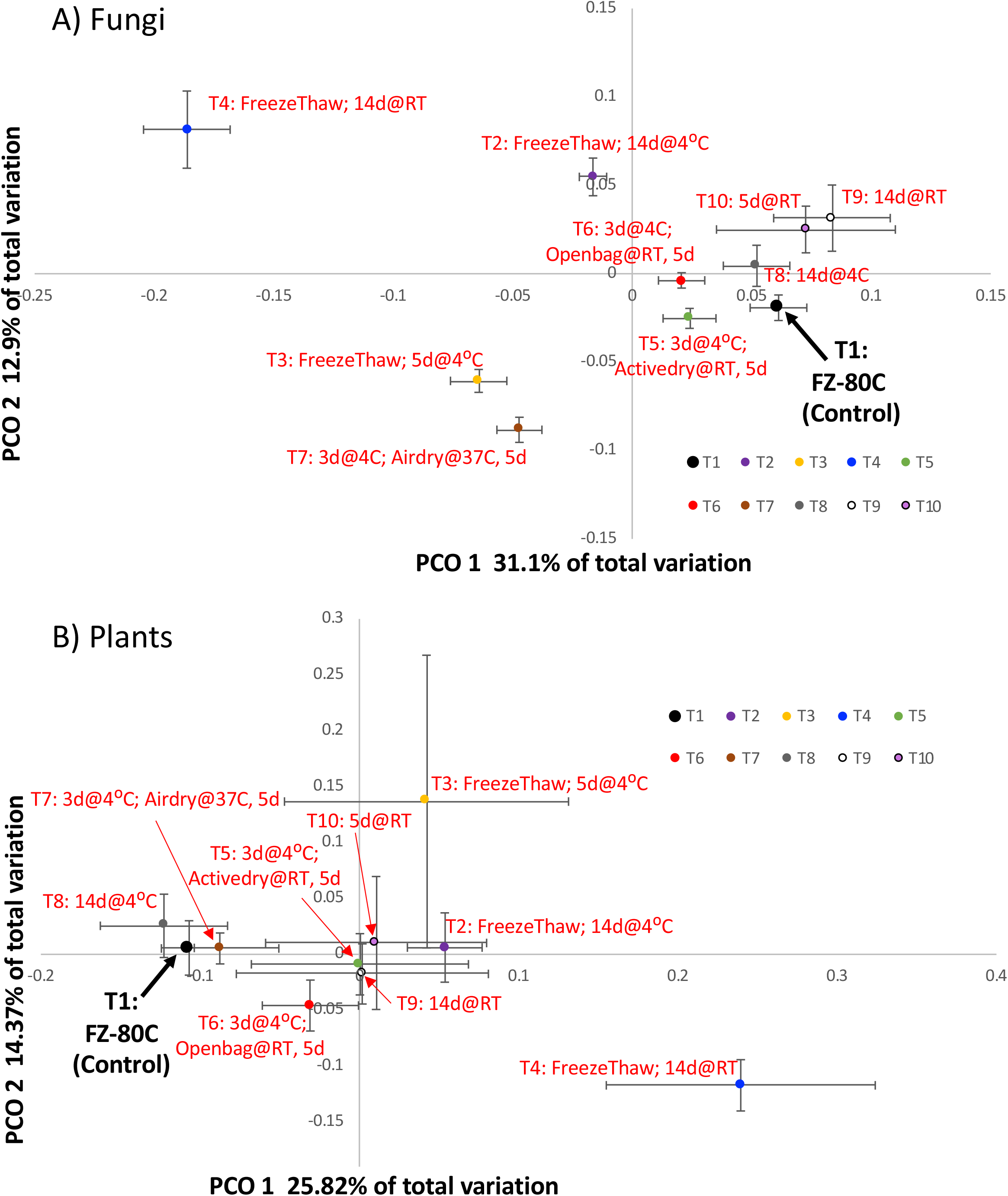
Principal coordinate diagrams of the fungal community data (A) and plant community data (B) highlighting the difference in community between the different soil storage treatments. Points show the mean axis scores and error bars show standard error of the mean.

**Fig. 2.**
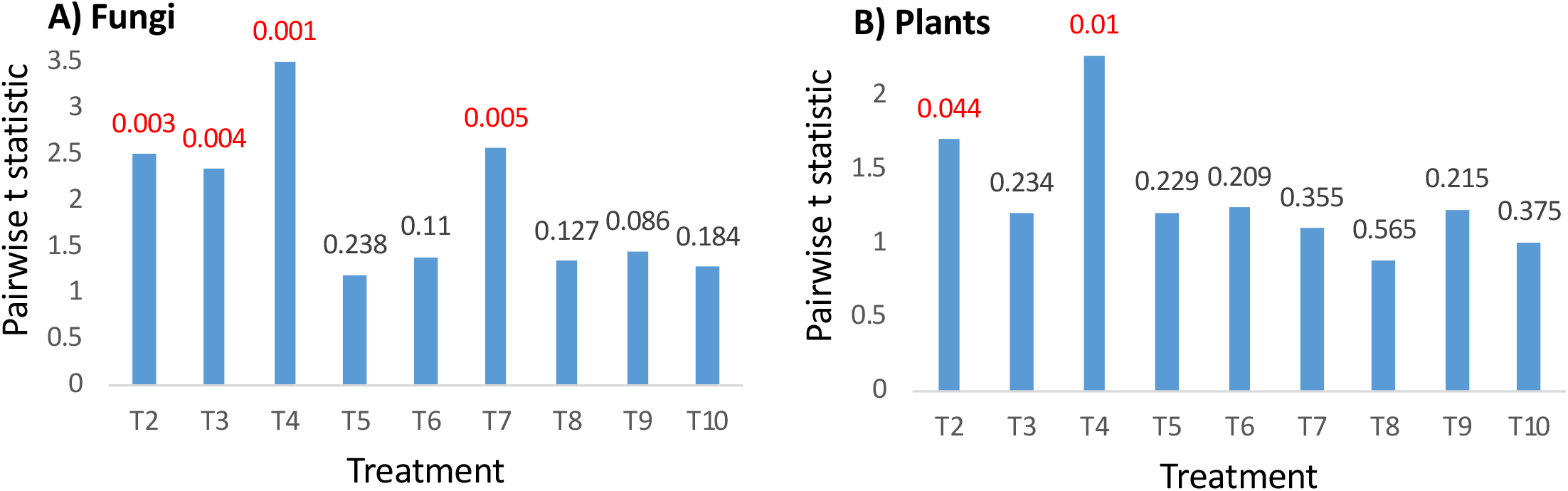
Levels of t-statistic from pairwise Permanova of the control treatment compared to all other treatments. The p value is shown above the bar with significant (P<0.05) values shown in red. (A) fungal community data (B) plant community data.

Similar analyses for the effect of different treatments on the plant DNA (including algae: Chlorophyta) remaining after storage show that the general level of divergence from the control was less than for fungi but still significant (Permanova Pseudo F = 2.5117 P= 0.001). Here too freeze-thaw treatments were also the most divergent (Fig. 1B, Fig. 2B) and there is similarity in the PCO ordinations for plant and fungal data. This similarity was corroborated by a Mantel test of the difference matrices, which revealed a Pearson correlation coefficient of 0.54 (P=0.0001). In contrast to the broader trend, the warm air-drying treatment (T7) had very different effect on the trajectory of the resulting plant and fungal communities later detected, causing significant change to the fungal community present (Fig. 2A) but very little effect on the plant DNA later recovered.

Apart from the treatments mentioned above, most treatments involving storage of soil at 4°C or at ambient temperature for up to 14 d did not result in significant changes to the plant or fungal populations later detected. In PCO ordination, the 4°C for 14 d (T8) treatment was closest to the control for both plants and fungi (Figs. 1A/1B). For the fungal community data, treatment 10 (closed bag, ambient temp) in particular showed an increase in the spread of data in both primary axis dimensions. For plants, in contrast to the fungi, the variance of axis scores (especially PCO1) for most treatments were larger than for the control treatment(Fig. 1B). This reduces the ability of statistical analyses to find a significant effect between treatments and may be related to the proliferation of spoilage fungi during storage.

Apart from green algae (Chlorophyta) which comprised <1% of total plant DNA in most treatments, the plant DNA present in the sieved soils was mainly within dead or dying tissues (e.g. fine roots). In contrast, a significant component of the fungal community would likely remain viable in the short term, with some species proliferating if storage conditions are conducive to their growth. Since fungi are the main decomposers of plant-derived lignocellulose in soil in terrestrial ecosystems, it is likely that proliferation of certain fungi would be associated with more rapid degradation of plant DNA. The relative sequence abundance of fungi and plants did significantly vary between treatments (Fig. 3). There were significant increases in the representation of fungi (and thus decrease in plants) relative to the control (T1), which was higher in two of the freeze-thaw treatments (T2 and T4) and T9 (23°C for 14 d). The relative proportions of fungi and plant sequences (Fig. 3) is clearly determined by the the exact mix of plant and fungal primers used but in our experiments the same mix was used for all samples, and therefore the changes reflect actual patterns of DNA degradation/proliferation.

**Fig. 3.**
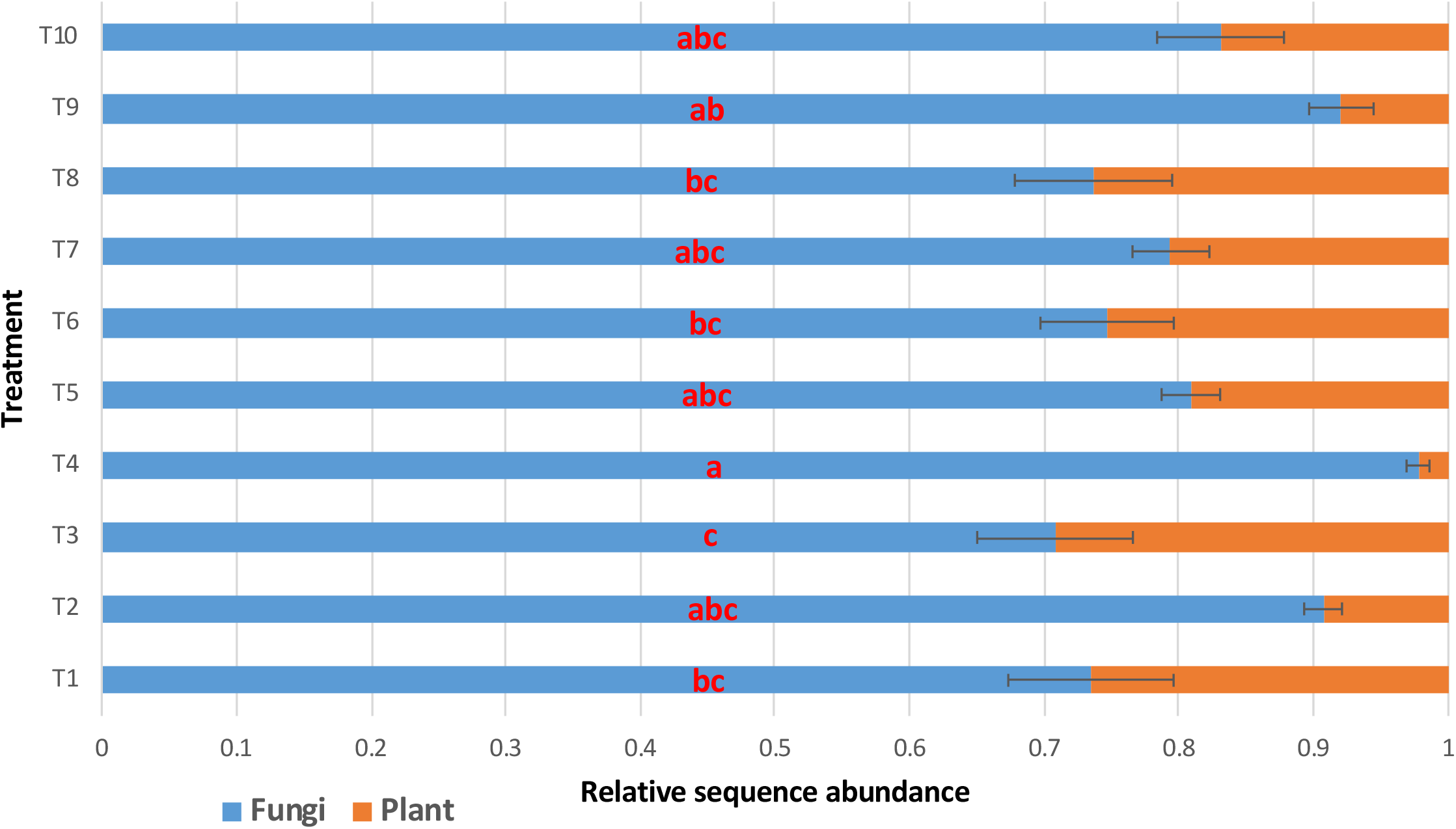
Relative sequence abundance of fungi to plants by treatment. Letters on the bars indicate significant groupings as determined by Tukey’s HSD post hoc test and error bars show standard error of the mean.

Some storage treatments led to a reduction in fungal species diversity (SuppData 3A/3B) relative to control, notably the freeze-thaw treatments (T2, T3, T4) and T7 (warm air-drying). In addition to significant differences in diversity between treatments, some treatments notably T5, T7 and T10, showed an increased spread of index values by replicate within treatment, as can be seen by the larger error bars. For plants (SuppData 3C/3D) reductions in species diversity were less pronounced, with only T4 differing significantly from control.

**SuppData 3**. Variations in diversity indices by storage treatment for the upland (Brignant) soil A) Fungi Simpson diversity index B) Fungi Shannon diversity index. C) Plant Simpson diversity index D) Plant Shannon diversity index. Letters above the bars indicate significant groupings as determined by Tukey’s HSD post hoc test and error bars show standard error of the mean. Note that Shannon and Simpson indices are scaled inversely (i.e. higher index = lower diversity).

More detailed examination of the differences in the fungal community composition with treatment reveal a large increase in abundance of Ascomycota relative to Basidiomycota for treatment 4 (Mean 2.83) compared to the control treatment (Mean 0.70) with all other treatments remaining very similar to the control (Fig. 4A). This change was mainly due to the ca. 10-fold (19.4 vs. 1.5%) increased abundance of the ascomycete *Metarhizium* (formerly Paecilomyces) *carneum* (UNITE species hypothesis SH1552520.08FU) following freeze-thaw and storage for 14d at 23°C (Fig. 4B). Mortierellomycota showed a similar 8-fold increase relative to control (17.8 vs 2.2%) in the freeze-thaw treatment stored at 4°C for 14 d and were also more abundant in treatments T4 and T8 (Fig. 4C).

**Fig. 4.**
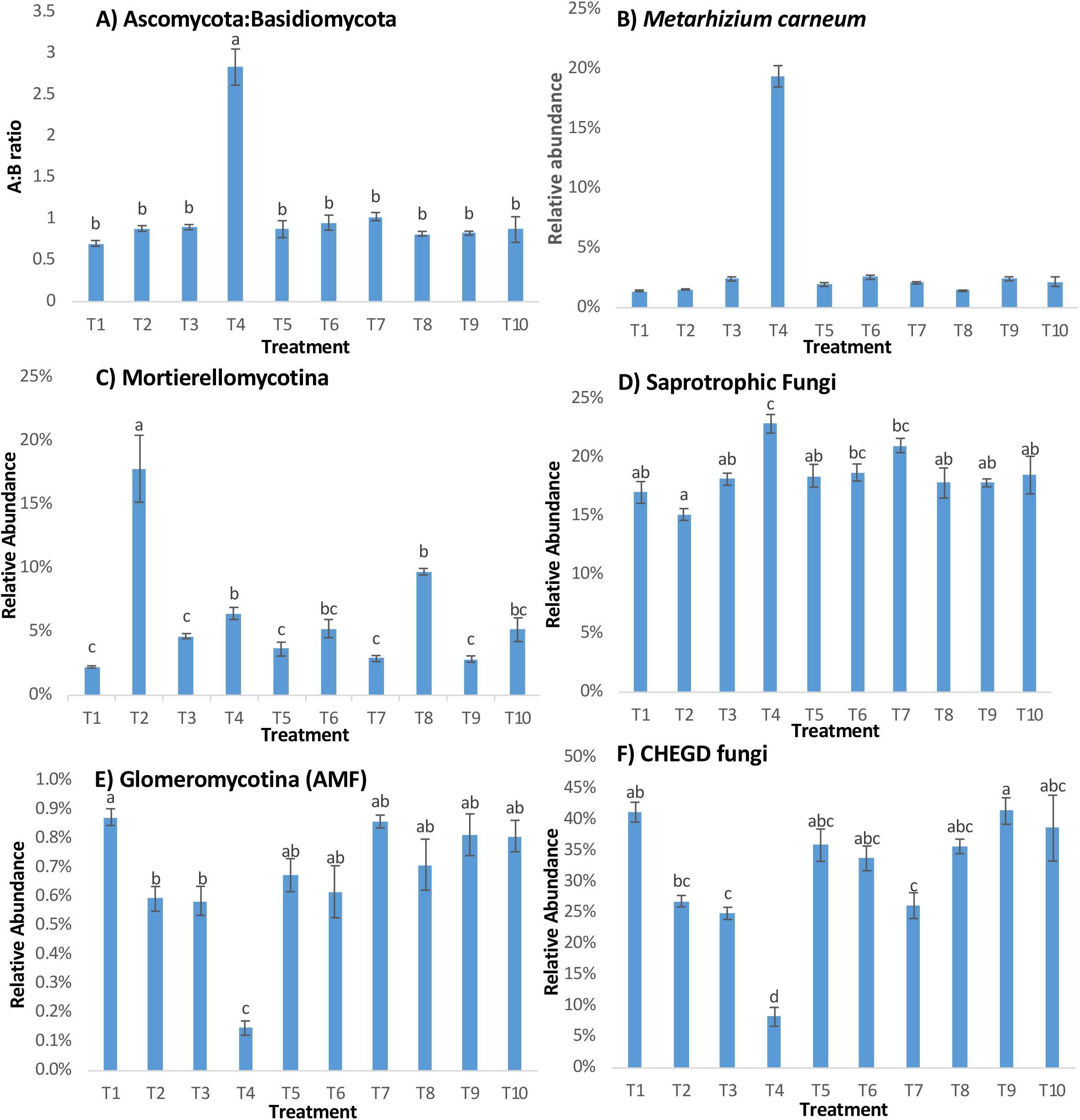
Variations in relative abundance of key fungal groups by storage treatment A) Ratio of Ascomycota to Basidiomycota; B) *Metarhizium carneum*; C) Mortierellomycotina; D) Saprotrophic fungi; E) Glomeromycotina (Arbuscular mycorrhizal fungi); F) Grassland fungi (CHEGD). Letters above the bars indicate significant groupings as determined by Tukey’s HSD post hoc test and error bars show standard error of the mean.

Analysis of the functional grouping, as determined unambiguously by FUNguild (Nguyen et al., 2016), revealed that ‘saprotrophic fungi’ demonstrated a similar but less pronounced trend, with a higher relative abundance in T4 relative to control (22.85% vs. 17.01%) (Fig. 4D). It should be noted that Mortierellomycota and *Metarhizium* spp. are classed as ‘symbiotroph’ and ‘animal pathogen’ respectively in FUNGuild.

As might be expected following disruption of active plant hosts, abundance of arbuscular mycorrhizal fungi (AMF; subphylum Glomeromycotina) was reduced under the three freeze-thaw storage conditions, with a 6-fold reduction in T4. Other fungi suspected to be mycorrhizal or with intricate biotrophic association with higher plants also showed large reductions in abundance, notably the CHEGD fungi. These fungi, mainly basidiomycetes and comprising members of the families <u>C</u>lavariaceae, <u>H</u>ygrophoraceae, <u>E</u>ntolomataceae and <u>G</u>eoglossaceae, are dominant components of undisturbed grassland habitats. Combined abundance of CHEGD fungi was 5-fold lower in treatment T4 and also significantly lower in treatments T2, T3 and T7 (Fig. 4D). Analysis of the individual components of the CHEG fungi revealed that the Clavariaceae, Hygrophoraceae, Geoglossomycetes varied by treatment with significantly lower relative abundances in treatment T4. However, there was no significant difference by treatment for Entolomataceae (SuppData 4). The two dominant CHEGD species in the original Brignant soil were *Clavulinopsis laeticolor* (UNITE SH1611741.08FU) and *Hygrocybe chlorophana* (SH1546991.08FU) with mean abundance in the control (T1) soil of 21.5% and 9.4% respectively, with these levels being 4-fold and 15-fold lower in the most unfavourable storage regime (T4; freeze-thaw followed by 14d at 23°C).

**SuppData 4**. Relative abundance of CHEG fungi by storage treatment for upland (Brignant) soil A) Hygrophoraceae B) Clavariaceae. C) Geoglossomycetes D) Entolomataceae. Letters above the bars indicate significant groupings as determined by Tukey’s HSD post hoc test and error bars show standard error of the mean.

In the Brignant soil, grasses (Poaceae; 9 spp.) were dominant (mean 88.8% of plant sequences in control treatment T1) followed by Brassicaceae (*Cardamine pratensis*; 5.57%), Asteraceae (3 spp.; 2.58%) and *Trifolium repens* (Fabaceae; 1.33%), with algae (Chlorophyta) comprising 0.41% of the plant sequences in the control soil. The turf layer was removed during sample collection, so the higher plant tissues comprised mainly (live or dead) root tissues. Several species (e.g. *Crepis capillaris, Hypochaeris radicata, Ranunculus repens, Cerastium glomeratum*) were detected in three or fewer of the initial 40 sieved soil samples, probably due to heterogeneous distribution of larger pieces of taproot tissue. The abundance of Poaceae varied by treatment (Suppdata 5) but this variation was not significant because of the broad range of data in some treatment replicates (e.g. 3.7% to 68.2% in treatment 4). The greatest treatment effect on plant populations was the ca. 20-fold increase in abundance of Chlorophyta in T4, likely due to tolerance of these microbes to freezing and later proliferation inside the clear plastic bags when incubated under ambient indoor lighting.

**Suppdata 5**. Relative sequence abundance of most abundant plant orders. Error bars show standard error of the mean.

### Alluvial (Gogerddan) soil

A subset of the storage treatments (T1, T3, T6, T8, T9) were applied to a contrasting soil type from an arable field in Gogerddan. The organic matter content of this soil was much lower (3.6% vs 7.3%) and initial plant and fungal populations of the original soils were very different. For example, Ascomycota fungi comprised ca. 70% of the initial fungal population at Gogerddan (vs 37% at Brignant), mostly due to the much lower abundance of CHEGD fungi (16% vs 41%, with Hygrophoraceae absent).

As with the Brignant soils, freeze-thaw storage (T3; freeze-thaw followed by 5 d at 23°C) resulted in the greatest difference in fungal populations relative to control (Fig. 5A), with a reduction in diversity (Suppdata 6A,6B) compared to control and the other treatments. The pattern of divergence of plant DNA composition followed a similar pattern to the fungi (Fig. 5B) but with no significant decrease in diversity indices (Suppdata 6C,6D). The most divergent treatment was storage at 23°C for 14 d (T9) which, in contrast with the other findings, showed higher diversity compared to the control but not the other treatments (Suppdata 6D).

**Fig. 5.**
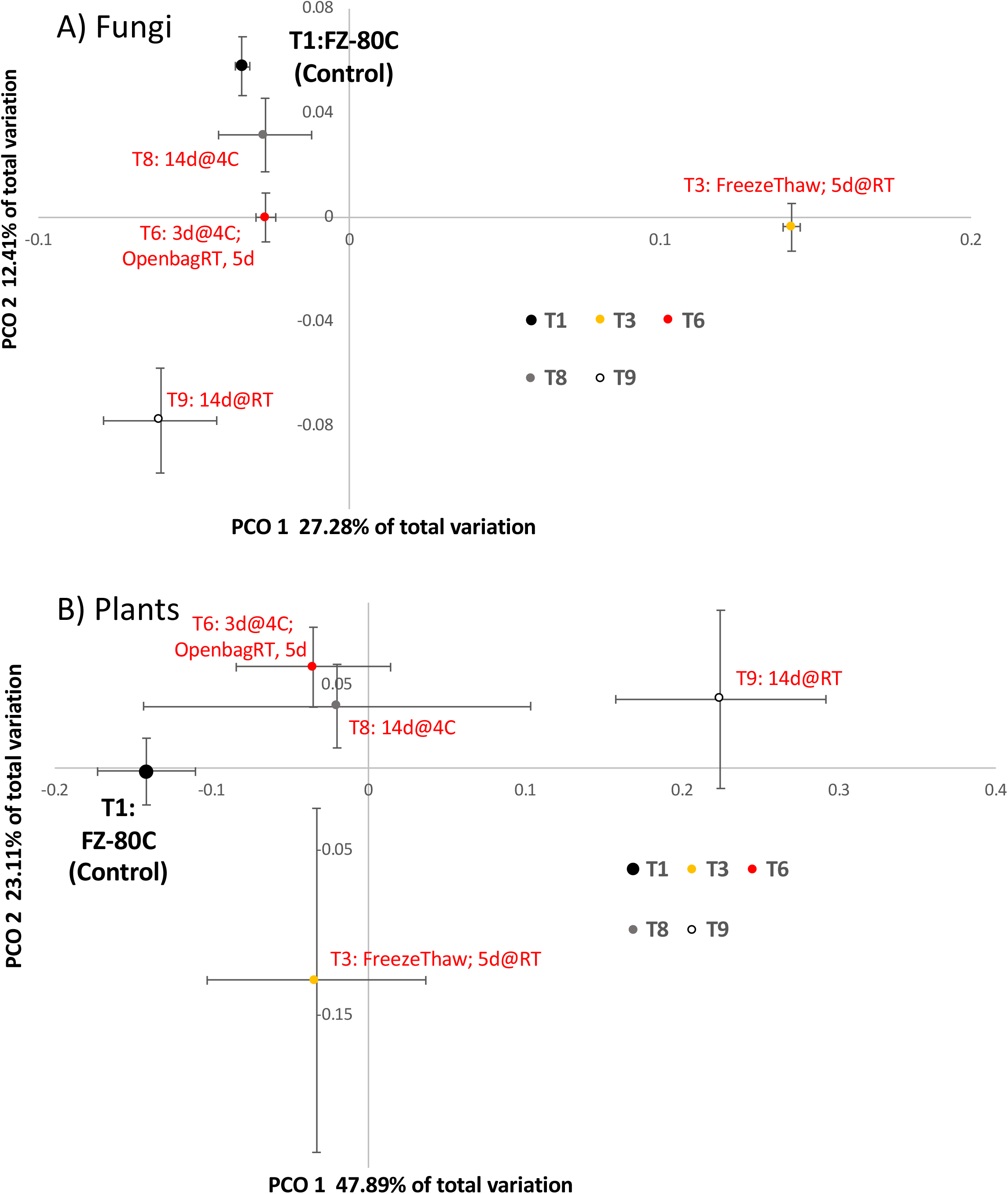
Principal coordinate diagrams of the fungal community data (A) and plant community data (B) in Gogerddan soil, highlighting the difference in community between the different soil storage treatments. Points show the mean axis scores and error bars indicate standard error of the mean. The control (immediate freezing at −80°C) is indicated with black arrow.

**SuppData 6**. Variations in diversity indices by storage treatment for the alluvial (Gogerddan) soil A) Fungi Simpson diversity index B) Fungi Shannon diversity index. C) Plant Simpson diversity index D) Plant Shannon diversity index. Letters above the bars indicate significant groupings as determined by Tukey’s HSD post hoc test and error bars show standard error of the mean. NS indicates no significant treatment effect. Note that Shannon and Simpson indices are scaled inversely (i.e. higher index = lower diversity).

In contrast to the Brignant soil, the relative abundance of Ascomycota:Basidiomycota did not increase following freeze-thaw storage (Suppdata 7), in large part because the dominant ascomycete at Gogerddan (Chaetothyriales_sp:SH1512803.08FU accounting for 21% of all fungal sequences in T1) was 4-fold lower in T3 and several basidiomycetous soil yeasts (e.g. *Solicoccozyma* spp.) increased several fold in abundance. In both the Gogerddan and Brignant soils, mycorrhizal fungi (AMF) declined in abundance following freeze-thaw treatment and fungi categorised as saprotrophic in FUNguild exhibited a 2-fold increase relative to control. *Metarhizium carneum* and Mortierellomycota (mainly *Mortierella* elongata) both increased 2-fold in abundance following freeze-thaw storage, as was found with the Brignant soil.

**SuppData 7**. Relative abundance of fungal groups by storage treatment for the alluvial soil (Gogerddan) A) Ascomycota to Basidiomycota ratio; B) *Metarhizium carneum*; C) Mortierellomycota; D) Saprotrophic fungi; E) Glomeromycotina (Arbuscular mycorrhizal fungi); F) Grassland fungi (CHEGD). Letters above the bars indicate significant groupings as determined by Tukey’s HSD post hoc test and error bars show standard error of the mean.

The plant community was also less diverse in the alluvial soil, with *Ranunculus bulbosus* (36.9%), the chlorophyte *Coelastrella* sp. (22.4%), Holcus lanatus (7.4%) and *Polygonum aviculare* (5.7%) found as the dominant plant species. Plant sequences comprised 14%-42% of all the sequences retrieved suggesting a similar ratio of plant to fungal biomass to that found in the Brignant soil, although with a greater abundance of Chlorophyta. The dominant chlorophyte, Coelastrella sp, is a ubiquitous species found in many substrates, including soils, worldwide (Wang et al., 2019). As with the Brignant soil, plant community data increased in variation with treatment especially after freeze thaw (T3) and longer storage at 4°C (T8) and room temperature (T9) (Fig. 5). As also observed for the Brignant soil, all treatments showed greater variability in plant community than the control (Fig. 5B-error bars), suggesting that DNA degradation was occurring and that this was due to the treatment effects rather than any initial inter-replicate variability. Seed germination was observed in some samples stored at room temperature (T9) and this may have contributed to this variability.

## Discussion

In this investigation we have tested the effectiveness of different soil storage conditions in stabilising fungal and plant DNA prior to later storage (−80°C) and DNA extraction. This is a concern for soil ecologists, since transport from remote and field sites to research laboratories requires interim storage in transit. This may also be a concern where soil sampling is undertaken by third parties and require transport by mail or courier. For example, the authors recently studied the soils of endemic woodlands in St. Helena and transport of samples to Wales involved storage of the samples for up to 8 d at 4°C in sealed plastic bags (Detheridge et al., 2020). On other occasions we receive samples from third party collaborators who may send soil samples in batches with delays of up to a week between collection and final frozen storage.

The data presented here shows clearly that refrigerated storage for up to 14 d (T8) prior to frozen storage at −80°C has little effect on the fungal or plant DNA later extracted. In contrast, samples initially frozen but allowed to thaw show the most rapid deterioration, presumably due to initial ice-damage from freezing and subsequent enzymatic degradation of DNA.

Air-drying (sometimes with the aid of silica gel) is widely used in botanical fieldwork for preservation of plant tissues (Chase and Hills, 1991; Liston et al., 1990) and has been shown to be superior to other methods for many (Pyle and Adams, 1989) but not all (Thomson, 2002) species. For the preservation of plant DNA in soil, this method was also highly effective, more so with passive, warm (T7) than active ambient air-drying (T5) (Fig. 1B). However, warm (37°C) air-drying caused large changes to fungal communities, presumably due to degradation of the DNA of certain fungi, notably CHEGD fungi (Fig. 4F). Relative to other groups of biota, fungi are heat-sensitive and only a few species can grow at 37°C (Robert et al., 2015) and no such effect was apparent with ambient air-drying. Mean relative abundance of saprotrophic fungi was greater for both air-dry treatments than in the control (Fig. 4D) but there was no significant treatment effect.

A few fungi were observed to increase in abundance under some storage treatments, presumably because the storage conditions were conducive to their growth. *Metarhizium* (formerly *Paecilomyces*) *carneum* (Kepler et al., 2014) showing the greatest increase in abundance, notably in the freeze-thawed soils which were then incubated at ambient temperature (Fig. 4C). This species is strongly chitinolytic and was frequently recovered from soil baited with chitin (Gray and Baxby, 1968; Jackson, 1965). Another closely related species, *Metarhizium marquandii* (Inglis and Tigano, 2006), showed the same patterns of relative abundance but was present at lower levels. Both these species are entomopathogenic to Lepidoptera (Bakeri et al., 2009; Magda and Said, 2014; Shin et al., 2013). However, in this experiment the soils were sieved did not have a high content of soil fauna and it is likely that proliferation of *M. carneum* was due to its ability to degrade the cell walls of recently dead fungi (e.g CHEGD fungi and AMF).

Of the 15 *Mortierella* spp. detected within the Brignant soil, all but one increased in abundance in the freeze-thawed soil incubated at 4°C (Fig. 4C). *Mortierella* spp. are psychro-tolerant (Melo et al., 2014; Widden, 1987), exhibit ice nucleation activity (Fröhlich-Nowoisky et al., 2015) and are abundant in recently thawed glacier forefront soils (Dresch et al., 2019). As has been found for *Metarhizium* spp., *Mortierella* spp. are also chitinolytic and frequently isolated in soil baiting experiments with chitin (Gray and Baxby, 1968; Jackson, 1965). *Mortierella alpina*, the most abundant *Mortierella* species found here is also reported to be parasitic on soil fungi (*Rhizoctonia* spp.) and nematodes (Al-Shammari et al., 2013), as well as occurring as endophytes of plant roots (Bonfante, 2020). Thus, it is likely that, like M. *carneum*, the *Mortierella* spp. benefit from the increased abundance dead hyphae (AMF/ CHEGD etc.) and are able to exploit these at low temperature. Mortierellomycotina (all *Mortierella* spp.) were also elevated 4-fold in soil stored at 4°C for 14 d (T8), the treatment least changed from the control. Thus the presence of elevated populations of these fungi provides a useful indication that soils have potentially been stored sub-optimally. Also highly elevated (10-fold) in freeze-thawed soils was *Myxotrixchum* (anamorph is Oidiodendron); members of this genus are also most commonly encountered in boreal soils rich in organic matter (Rice and Currah, 2005).

Of the taxa which declined substantially following freeze-thawing, Glomeromycotina (AMF) showed the greatest decline. However, within this subphylum some taxa were more heavily affected than others. For instance, Acaulospora sp. showed a >4-fold decline (treatments T2 and T4), whereas Claroideoglomus spp. declined less than 2-fold. This is consistent with the findings of Klironomos et al. (2001) who found Claroideoglomus to be tolerant of freeze-thaw cycles compared to other AMF spp.

The CHEGD fungi (barring Entolomataceae) also showed substantial decline in relative abundance in freeze-thaw treatments. Like AMF, these fungi are obligate root-associated biotrophs (Halbwachs et al., 2013; Halbwachs et al., 2018), and are negatively affected by killing of host vegetation (Griffith et al., 2014). the fact that Entolomataceae were differently affected to other CHEGD fungi suggests that they are nutritionally more flexible, potentially with some saprotrophic ability. Together the CHEGD fungi comprised >40% of the total fungal biomass at the Brignant site and are recognized to be the dominant fungi of undisturbed mesotrophic grasslands (Griffith et al., 2019; Halbwachs et al., 2013). Their susceptibility to freeze-thaw treatment and resultant increase in fungal necromass is likely the cause of the large proliferation of the chitinolytic M. *carneum* and *Mortierella* spp. in freeze-thaw treatments.

## Conclusions

To our knowledge this is the only study to have examined the effects of sub-optimal soil storage on eukaryotic eDNA using a high resolution method. When analysing fungal communities, for those situations where freezing samples and freeze drying are impractical, such as remote locations without equipment and requiring length shipping times, the analysis indicates that the best options available are to ship cold or, if impractical, to air dry at room temperature prior to shipping. Air drying can be enhanced by using an unheated active air source, such as a blower or a fan. Pre-freezing a sample prior to shipping is not recommended, nor is drying with a heat source such as a drying oven. For two contrasting soil types, the results of suboptimal storage were similar, suggesting broad applicability of these guidelines.

## Acknowledgements

This was supported by a grant to GWG, APD, LAC and JS from Welsh Assembly Government (Contract C343/2017/2108; “Higher Plant DNA Sequencing in Soil” via David Rogerson and Geraint Lewis), and capacity developed as part of the Welsh European Funding Office Flexis West project C80835 (APD and JS). The Institute of Biological, Environmental, and Rural Sciences receives strategic funding from the BBSRC.

## Author’s contribution

GWG, JS, APD conceived the study. Experiments were undertaken by LAC and APD. Manuscript was drafted by GWG/APD and all authors contributed to editing of the manuscript.

## References

Abarenkov, K., Zirk, A., Piirmann, T., Pöhönen, R., Ivanov, F., Nilsson, R.H., Kõljalg, U., 2019. Full mothur UNITE+INSD dataset 1. Version 02.02.2019.

Al-Shammari, T.A., Bahkali, A.H., Elgorban, A.M., El-Kahky, M.T., Al-Sum, B.A., 2013. The use of Trichoderma longibrachiatum and *Mortierella* alpina against root-knot nematode, Meloidogyne javanica on tomato. J.ournal of Pure and Applied Microbiology 7, 199–207.

Bainard, L.D., Klironomos, J.N., Hart, M.M., 2010. Differential effect of sample preservation methods on plant and arbuscular mycorrhizal fungal DNA. Journal of Microbiological Methods 82, 124–130.

Bakeri, S.A., Ali, S.R.A., Tajuddin, N.S., Kamaruzzaman, N.E., 2009. Efficacy of entomopathogenic fungi, Paecilomyces spp., in controlling the oil palm bagworm, Pteroma pendula (Joannis). Journal of Oil Palm Research 21, 693–699.

Bonfante, P., 2020. Mucoromycota: going to the roots of plant-interacting fungi. Fungal Biology Reviews.

Castaño, C., Parladé, J., Pera, J., De Aragón, J.M., Alday, J.G., Bonet, J.A., 2016. Soil drying procedure affects the DNA quantification of Lactarius vinosus but does not change the fungal community composition. Mycorrhiza 26, 799–808.

Clark, I.M., Hirsch, P.R., 2008. Survival of bacterial DNA and culturable bacteria in archived soils from the Rothamsted Broadbalk experiment. Soil Biology and Biochemistry 40, 1090–1102.

Chase, M.W., Hills, H.H., 1991. Silica gel: an ideal material for field preservation of leaf samples for DNA studies. Taxon 40, 215–220.

Chen, S., Yao, H., Han, J., Liu, C., Song, J., Shi, L., Zhu, Y., Ma, X., Gao, T., Pang, X., Luo, K., Li, Y., Li, X., Jia, X., Lin, Y., Leon, C., 2010. Validation of the ITS2 Region as a Novel DNA Barcode for Identifying Medicinal Plant Species. PLOS ONE 5, e8613.

Detheridge, A.P., Brand, G., Fychan, R., Crotty, F.V., Sanderson, R., Griffith, G.W., Marley, C.L., 2016. The legacy effect of cover crops on soil fungal populations in a cereal rotation. Agriculture, Ecosystems & Environment 228, 49–61.

Detheridge, A.P., Comont, D., Callaghan, T.M., Bussell, J., Brand, G., Gwynn-Jones, D., Scullion, J., Griffith, G.W., 2018. Vegetation and edaphic factors influence rapid establishment of distinct fungal communities on former coal-spoil sites. Fungal Ecology 33, 92–103.

Detheridge, A.P., Cherrett, S., Clasen, L.A., Medcalf, K., Pike, S., Griffith, G.W., Scullion, J., 2020. Depauperate soil fungal populations from the St. Helena endemic Commidendrum robustum are dominated by Capnodiales. Fungal Ecology 45, 100911.

Dresch, P., Falbesoner, J., Ennemoser, C., Hittorf, M., Kuhnert, R., Peintner, U., 2019. Emerging from the ice-fungal communities are diverse and dynamic in earliest soil developmental stages of a receding glacier. Environmental Microbiology 21, 1864–1880.

Epp, L.S., Boessenkool, S., Bellemain, E.P., Haile, J., Esposito, A., Riaz, T., Erseus, C., Gusarov, V.I., Edwards, M.E., Johnsen, A., 2012. New environmental metabarcodes for analysing soil DNA: potential for studying past and present ecosystems. Molecular Ecology 21, 1821–1833.

Fröhlich-Nowoisky, J., Hill, T.C., Pummer, B.G., Yordanova, P., Franc, G.D., Pöschl, U., 2015. Ice nucleation activity in the widespread soil fungus *Mortierella* alpina. Biogeosciences 12, 1057–1071.

Geml, J., Gravendeel, B., van der Gaag, K.J., Neilen, M., Lammers, Y., Raes, N., Semenova, T.A., de Knijff, P., Noordeloos, M.E., 2014. The contribution of DNA metabarcoding to fungal conservation: Diversity assessment, habitat partitioning and mapping Red-Listed fungi in protected coastal Salix repens communities in the Netherlands. PLOS ONE 9, e99852.

Gray, T., Baxby, P., 1968. Chitin decomposition in soil: II. The ecology of chitinoclastic micro-organisms in forest soil. Transactions of the British Mycological Society 51, 293–309.

Griffith, G.W., Cavalli, O., Detheridge, A.P., 2019. An assessment of the fungal conservation value of Hardcastle Crags using NextGen DNA sequencing (NECR258). Natural England Commissioned Report NECR258 (http://publications.naturalengland.org.uk/publication/4543317115404288).

Griffith, G.W., Graham, A., Woods, R.G., Easton, G.L., Halbwachs, H., 2014. Effect of biocides on the fruiting of waxcap fungi. Fungal Ecology 7, 67–69.

Nguyen, N.H., Song, Z., Bates, S.T., Branco, S., Tedersoo, L., Menke, J., Schilling, J.S., Kennedy, P.G., 2016. FUNGuild: an open annotation tool for parsing fungal community datasets by ecological guild. Fungal Ecology 20, 241–248.

Halbwachs, H., Dentinger, B.T.M., Detheridge, A.P., Karasch, P., Griffith, G.W., 2013. Hyphae of waxcap fungi colonise plant roots. Fungal Ecology 6, 487–492.

Halbwachs, H., Easton, G.L., Bol, R., Hobbie, E.A., Garnett, M.H., Peršoh, D., Dixon, L., Ostle, N., Karasch, P., Griffith, G.W., 2018. Isotopic evidence of biotrophy and unusual nitrogen nutrition in soil-dwelling Hygrophoraceae. Environmental Microbiology 20, 3573–3588.

Hallett, S.H., Sakrabani, R., Keay, C., Hannam, J.A., 2017. Developments in land information systems: examples demonstrating land resource management capabilities and options. Soil Use and Management 33, 514–529.

Inglis, P.W., Tigano, M.S., 2006. Identification and taxonomy of some entomopathogenic Paecilomyces spp.(Ascomycota) isolates using rDNA-ITS sequences. Genetics and Molecular Biology 29, 132–136.

ISO-11063, 2012. Soil quality — Method to directly extract DNA from soil samples. International Organization for Standardizaltion (ISO). https://www.iso.org/obp/ui/#iso:std:iso:11063:ed-1:v1:en.

ISO-18400-206, 2002. Soil quality — Sampling — Part 206: Collection, handling and storage of soil under aerobic conditions for the assessment of microbiological processes, biomass and diversity in the laboratory. International Organization for Standardizaltion (ISO). https://www.iso.org/standard/68249.html.

Jackson, R., 1965. Studies of fungi in pasture soils: III. Physiological studies on some fungal isolates from the root surface and from organic debris. New Zealand Journal of Agricultural Research 8, 878–888.

Keepers, K.G., Pogoda, C.S., White, K.H., Anderson Stewart, C.R., Hoffman, J.M., Ruiz, A.M., McCain, C.M., Lendemer, J.C., Kane, N.C., Tripp, E.A., 2019. Whole genome shotgun sequencing detects greater lichen fungal diversity than amplicon-based methods in environmental samples. Frontiers in Ecology and Evolution 7, 484.

Kennedy, N.A., Walker, A.W., Berry, S.H., Duncan, S.H., Farquarson, F.M., Louis, P., Thomson, J.M., 2014. The impact of different DNA extraction kits and laboratories upon the assessment of human gut microbiota composition by 16S rRNA gene sequencing. PloS one 9.

Kepler, R.M., Humber, R.A., Bischoff, J.F., Rehner, S.A., 2014. Clarification of generic and species boundaries for *Metarhizium* and related fungi through multigene phylogenetics. Mycologia 106, 811–829.

Klironomos, J.N., Hart, M.M., Gurney, J.E., Moutoglis, P., 2001. Interspecific differences in the tolerance of arbuscular mycorrhizal fungi to freezing and drying. Canadian Journal of Botany 79, 1161–1166.

Latch, E.K., 2020. Integrating genomics into conservation management. Molecular Ecology Resources 20, doi: 10.1111/1755-0998.13188.

Lauber, C.L., Zhou, N., Gordon, J.I., Knight, R., Fierer, N., 2010. Effect of storage conditions on the assessment of bacterial community structure in soil and human-associated samples. FEMS Microbiology Letters 307, 80–86.

Lee, Y.B., Lorenz, N., Dick, L.K., Dick, R.P., 2007. Cold storage and pretreatment incubation effects on soil microbial properties. Soil Science Society of America Journal 71, 1299–1305.

Lindahl, B.D., Nilsson, R.H., Tedersoo, L., Abarenkov, K., Carlsen, T., Kjøller, R., Kõljalg, U., Pennanen, T., Rosendahl, S., Stenlid, J., 2013. Fungal community analysis by high-throughput sequencing of amplified markers–a user’s guide. New Phytologist 199, 288–299.

Liston, A., Rieseberg, L.H., Adams, R.P., Do, N., Ge-lin, Z., 1990. A method for collecting dried plant specimens for DNA and isozyme analyses, and the results of a field test in Xinjiang, China. Annals of the Missouri Botanical Garden 77, 859–863.

Magda, S., Said, S., 2014. Efficacy of two entomopathogenic fungi against corn pests under laboratory and field conditions in Egypt. European Journal of Academic Essays 1, 1–6.

Martí, E., Càliz, J., Montserrat, G., Garau, M.A., Cruañas, R., Vila, X., Sierra, J., 2012. Air-drying, cooling and freezing for soil sample storage affects the activity and the microbial communities from two Mediterranean soils. Geomicrobiology Journal 29, 151–160.

Melo, I.S., Santos, S.N., Rosa, L.H., Parma, M.M., Silva, L.J., Queiroz, S.C., Pellizari, V.H., 2014. Isolation and biological activities of an endophytic *Mortierella* alpina strain from the Antarctic moss Schistidium antarctici. Extremophiles 18, 15–23.

OECD, 2000. OECD Guidelines for the Testing Of Chemicals. Test No. 217: Soil Microorganisms: Carbon Transformation Test (doi.org/10.1787/20745761), Paris.

Ogwu, M.C., Kerfahi, D., Song, H., Dong, K., Seo, H., Lim, S., Srinivasan, S., Kim, M.K., Waldman, B., Adams, J.M., 2019. Changes in soil taxonomic and functional diversity resulting from gamma irradiation. Scientific Reports 9, 1–13.

Orgiazzi, A., Dunbar, M.B., Panagos, P., de Groot, G.A., Lemanceau, P., 2015. Soil biodiversity and DNA barcodes: opportunities and challenges. Soil Biology and Biochemistry 80, 244–250.

Petric, I., Philippot, L., Abbate, C., Bispo, A., Chesnot, T., Hallin, S., Laval, K., Lebeau, T., Lemanceau, P., Leyval, C., 2011. Inter-laboratory evaluation of the ISO standard 11063 “Soil quality—Method to directly extract DNA from soil samples”. Journal of Microbiological Methods 84, 454–460.

Pyle, M.M., Adams, R.P., 1989. In situ preservation of DNA in plant specimens. Taxon 38, 576–581.

R_Core_Team, 2013. R: A language and environment for statistical computing. R Foundation for Statistical Computing, Vienna, Austria (URL http://www.R-project.org/).

Rice, A.V., Currah, R.S., 2005. Oidiodendron: A survey of the named species and related anamorphs of Myxotrichum. Studies in Mycology 53, 83–120.

Robert, V., Cardinali, G., Casadevall, A., 2015. Distribution and impact of yeast thermal tolerance permissive for mammalian infection. BMC biology 13, 18.

Rubin, B.E., Gibbons, S.M., Kennedy, S., Hampton-Marcell, J., Owens, S., Gilbert, J.A., 2013. Investigating the impact of storage conditions on microbial community composition in soil samples. PloS One 8, e70460.

Shin, T.Y., Lee, W.W., Ko, S.H., Choi, J.B., Bae, S.M., Choi, J.Y., Lee, K.S., Je, Y.H., Jin, B.R., Woo, S.D., 2013. Distribution and characterisation of entomopathogenic fungi from Korean soils. Biocontrol Science and Technology 23, 288–304.

Soliman, T., Yang, S.-Y., Yamazaki, T., Jenke-Kodama, H., 2017. Profiling soil microbial communities with next-generation sequencing: the influence of DNA kit selection and technician technical expertise. PeerJ 5, e4178.

Straube, D., Juen, A., 2013. Storage and shipping of tissue samples for DNA analyses: A case study on earthworms. European Journal of Soil Biology 57, 13–18.

Taberlet, P., Prud’homme, S.M., Campione, E., Roy, J., Miquel, C., Shehzad, W., Gielly, L., Rioux, D., Choler, P., Clément, J.-C., 2012. Soil sampling and isolation of extracellular DNA from large amount of starting material suitable for metabarcoding studies. Molecular Ecology 21, 1816–1820.

Tedersoo, L., Bahram, M., Põlme, S., Kõljalg, U., Yorou, N.S., Wijesundera, R., Ruiz, L.V., Vasco-Palacios, A.M., Thu, P.Q., Suija, A., 2014. Global diversity and geography of soil fungi. Science 346, 1256688.

Terrat, S., Plassart, P., Bourgeois, E., Ferreira, S., Dequiedt, S., Adele-Dit-De-Renseville, N., Lemanceau, P., Bispo, A., Chabbi, A., Maron, P.A., 2015. Meta-barcoded evaluation of the ISO standard 11063 DNA extraction procedure to characterize soil bacterial and fungal community diversity and composition. Microbial Biotechnology 8, 131–142.

Thomson, J.A., 2002. An improved non-cryogenic transport and storage preservative facilitating DNA extraction from ‘difficult’plants collected at remote sites. Telopea 9, 755–760.

Valentin, R.E., Fonseca, D.M., Gable, S., Kyle, K.E., Hamilton, G.C., Nielsen, A.L., Lockwood, J.L., 2020. Moving eDNA surveys onto land: Strategies for active eDNA aggregation to detect invasive forest insects. Molecular Ecology Resources.

Wang, Q., Song, H., Liu, X., Liu, B., Hu, Z., Liu, G., 2019. Morphology and molecular phylogeny of coccoid green algae Coelastrella sensu lato (Scenedesmaceae, Sphaeropeales), including the description of three new species and two new varieties. Journal of Phycology 55, 1290–1305.

Weißbecker, C., Buscot, F., Wubet, T., 2017. Preservation of nucleic acids by freeze-drying for next generation sequencing analyses of soil microbial communities. Journal of Plant Ecology 10, 81–90.

Widden, P., 1987. Fungal communities in soils along an elevation gradient in northern England. Mycologia 79, 298–309.

Williams, N., 2020. Fungal DNA and conservation. Conservation Land Management 18, 9–15.

