## Supplementary material for "Soil stabilisatizion for DNA metabarcoding of plants and fungi. Implications for sampling at remote locations or via third-parties": SupplmentaryData

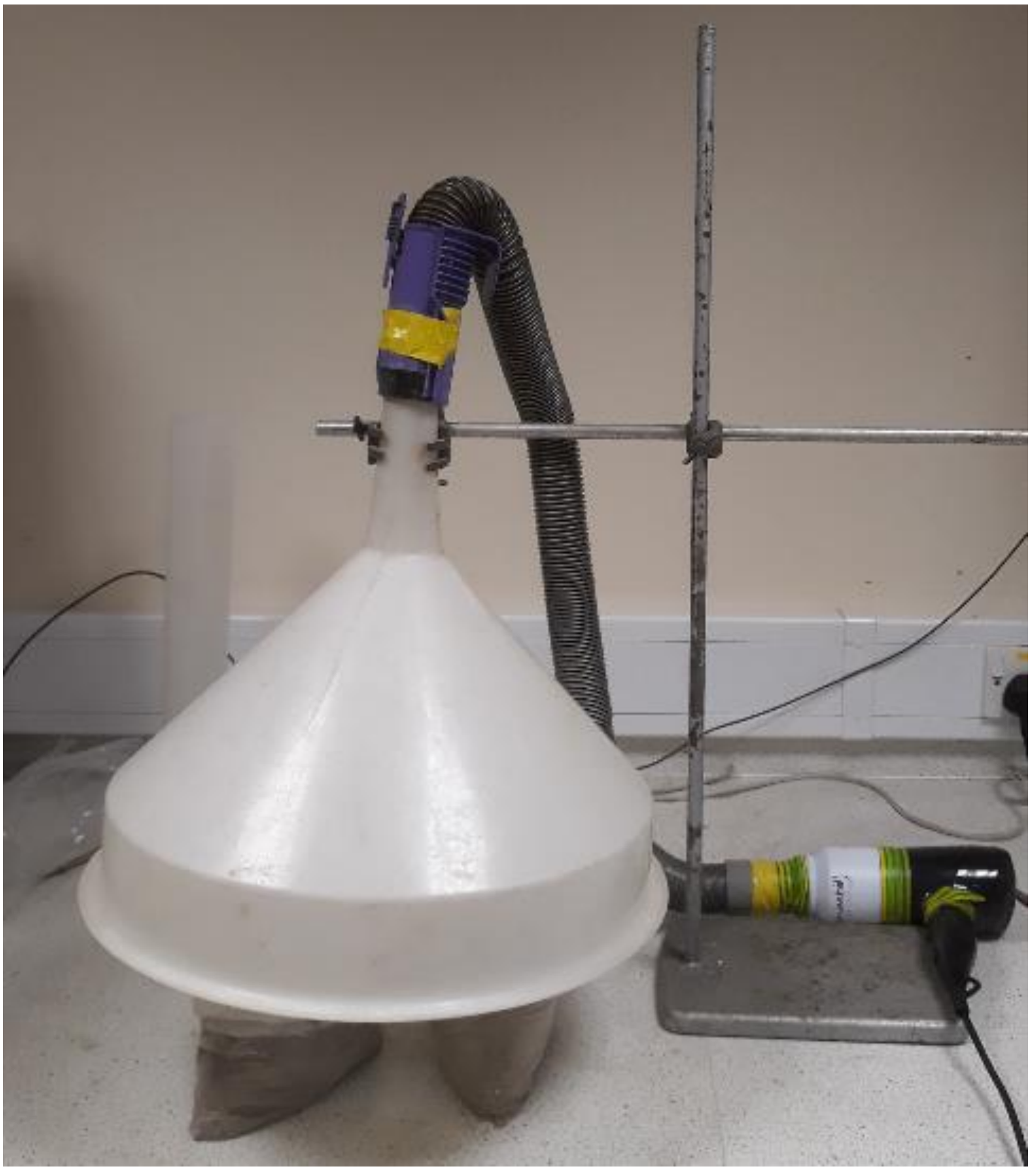

**SuppData 1.** Photo of the hairdrier apparatus used for active air-drying of soil (treatment T5)

**SuppData 2.** Fungal forward primer sequences with target group to amplify all fungal groups, and also Stramenopiles (Ooomyces) devised by Tedersoo et al. (2014)

| Name | Sequence | Target Group |
|---|---|---|
| ITS3NGS1 | CTAGACTCGTCATCGATGAAGAACGCAG | Ca. 95% of all fungi |
| ITS3NGS2 | CTAGACTCGTCAACGATGAAGAACGCAG | Chytridiomycota |
| ITS3NGS3 | CTAGACTCGTCACCGATGAAGAACGCAG | Sebacinales p.parte |
| ITS3NGS4 | CTAGACTCGTCATCGATGAAGAACGTAG | Glomeromycota |
| ITS3NGS5 | CTAGACTCGTCATCGATGAAGAACGTGG | Sordariales p.parte |
| ITS3NGS10 | CTAGACTCGTCATCGATGAAGAACGCTG | Stramenopila |

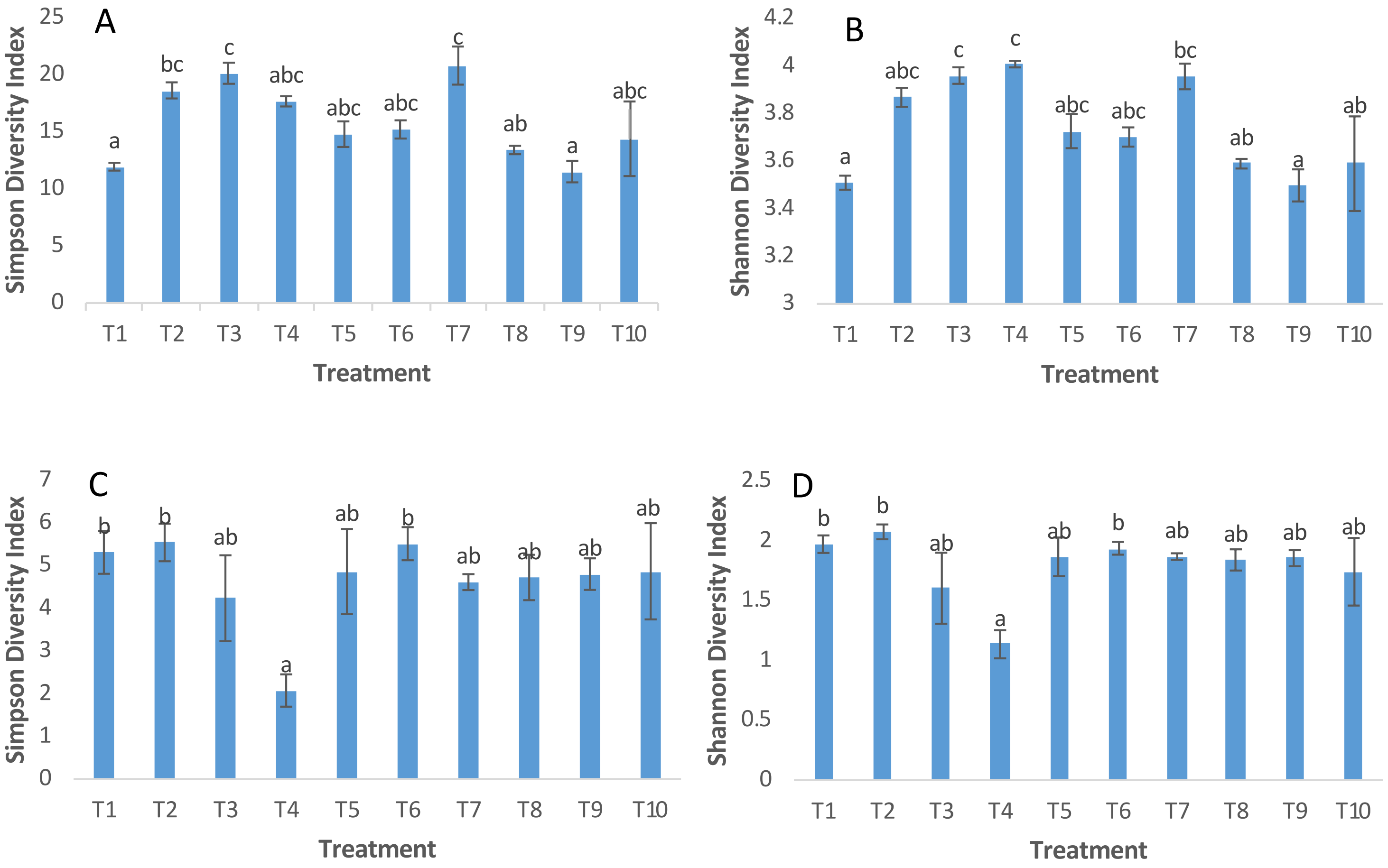


**SuppData 3.** Variations in diversity indices by storage treatment for the upland (Brignant) soil A) Fungi Simpson diversity index B) Fungi Shannon diversity index. C) Plant Simpson diversity index D) Plant Shannon diversity index. Letters above the bars indicate significant groupings as determined by Tukey's HSD post hoc test and error bars show standard error of the mean. Note that Shannon and Simpson indices are scaled inversely (i.e. higher index = lower diversity).

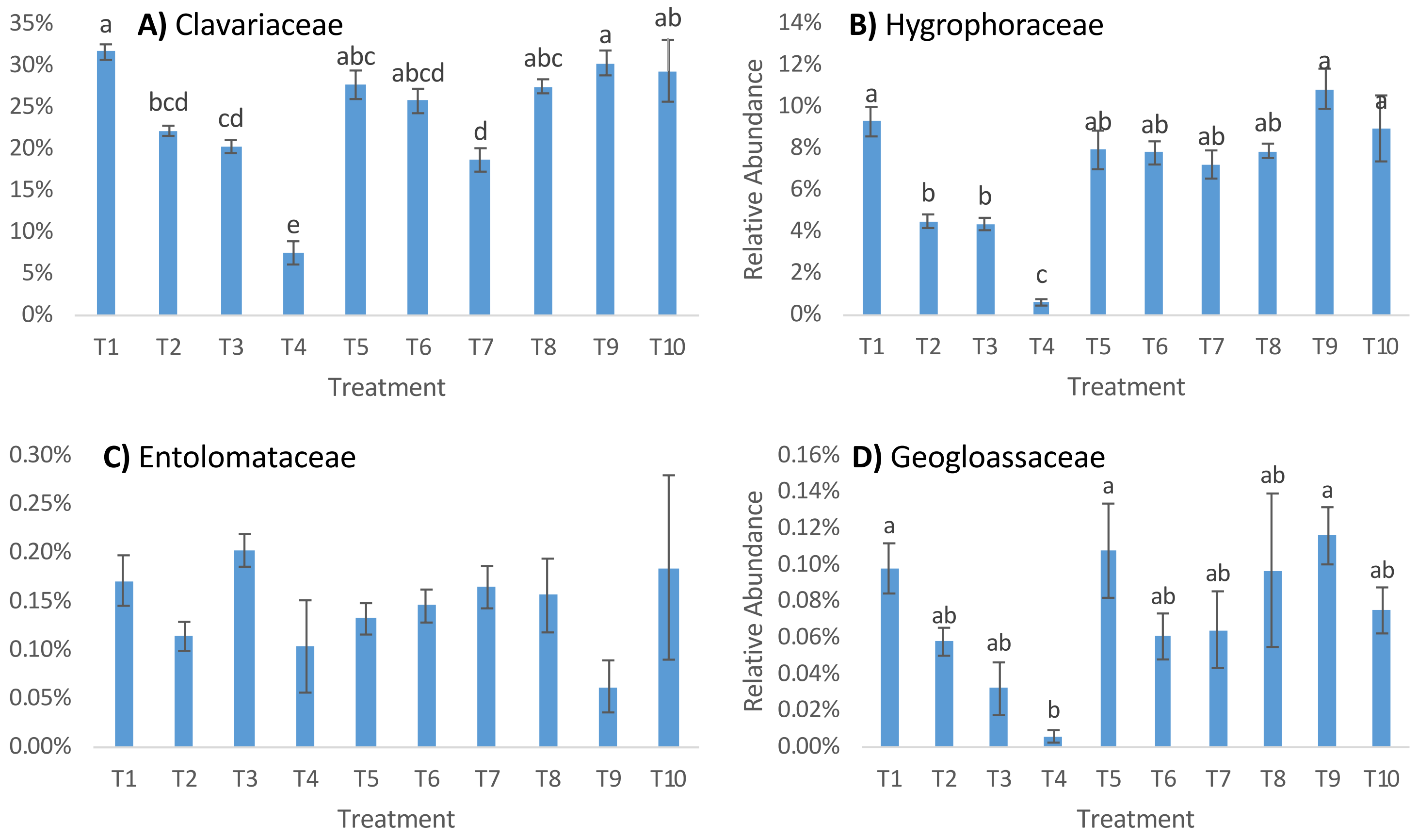


**SuppData 4.** Relative abundance of CHEG fungi by storage treatment for (Brignant soil). A) Clavariaceae; B) Hygrophoraceae; C) Entolomataceae; D) Geoglossaceae. Letters above the bars indicate significant groupings as determined by Tukey's HSD post hoc test and error bars show standard error of the mean.

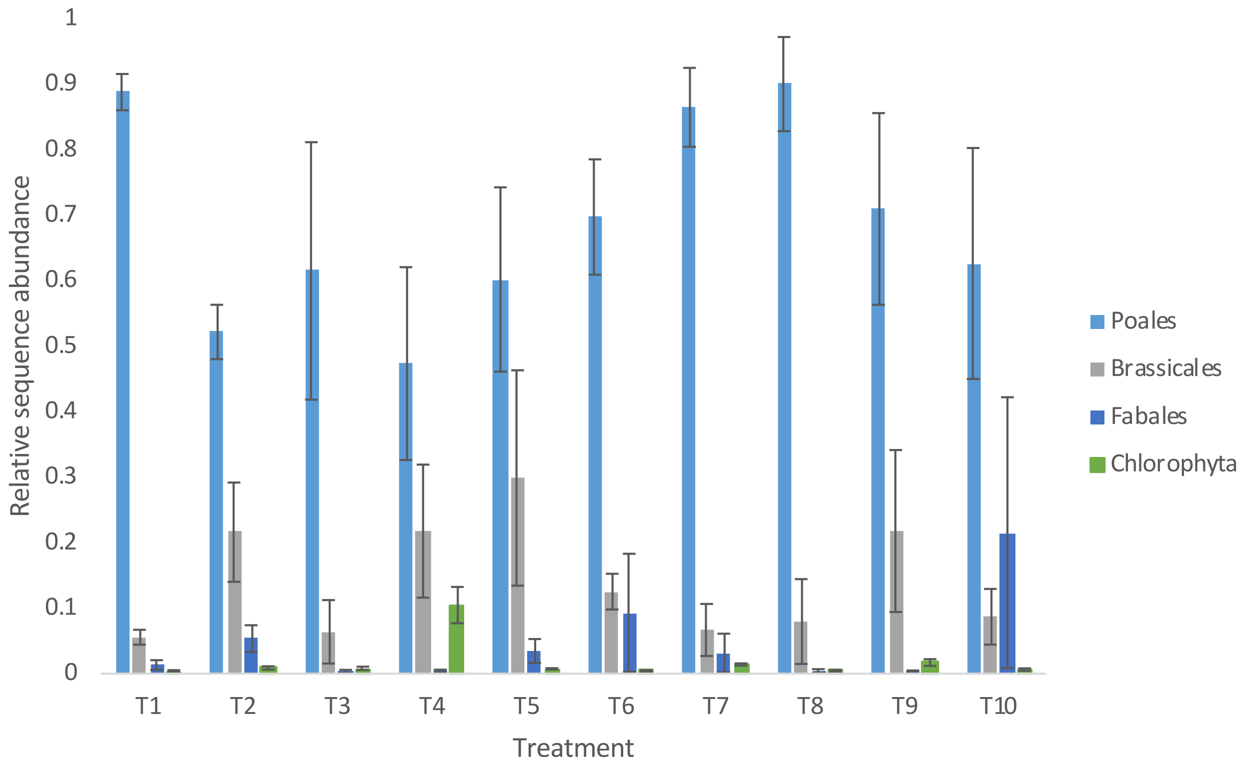


**Suppdata 5.** Relative sequence abundance of most abundant plant orders and Chlorophyta (Brignant soil). Error bars show standard error of the mean.

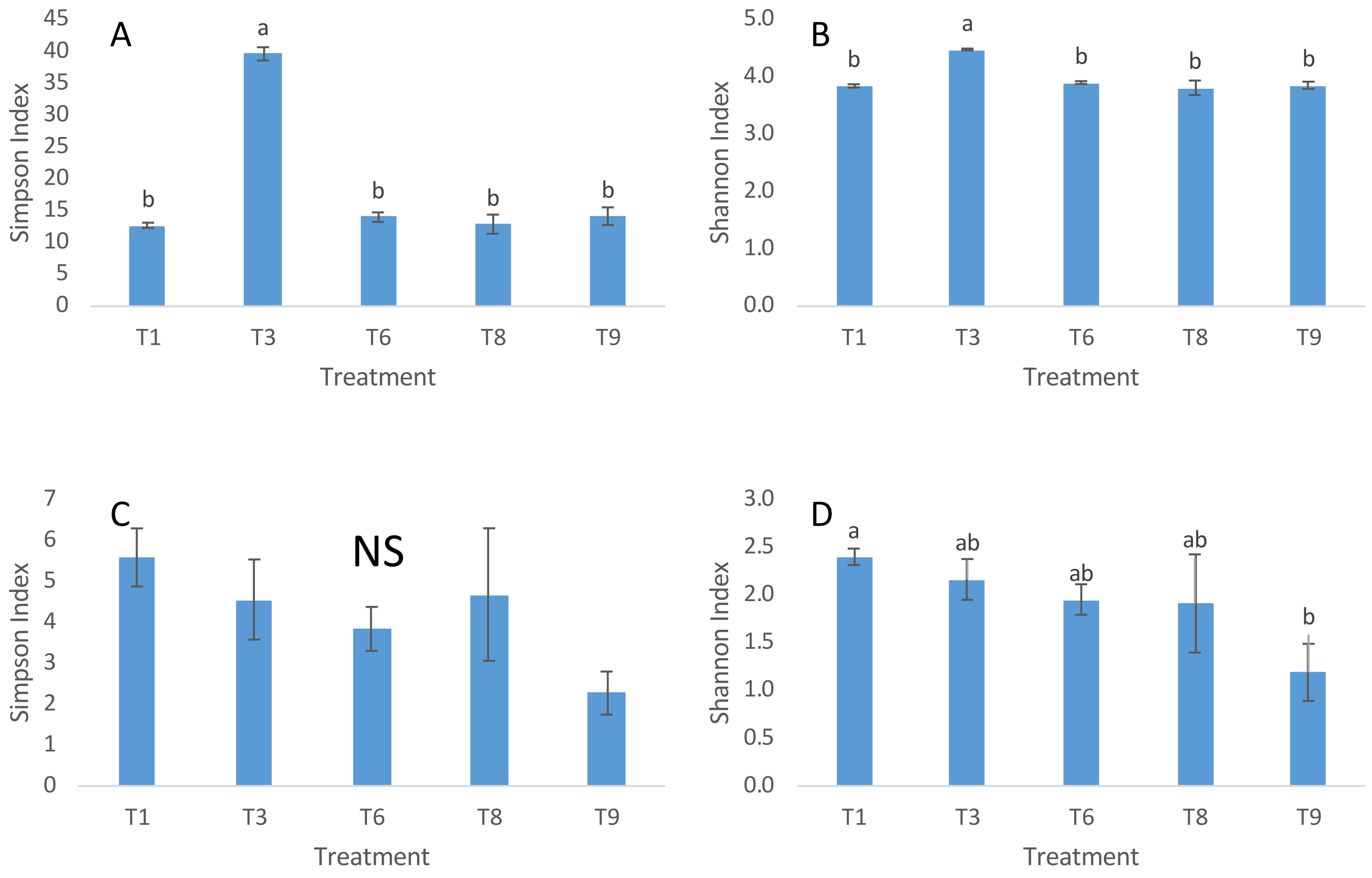


**SuppData 6.** Variations in diversity indices by storage treatment for the alluvial (Gogerddan) soil A) Fungi Simpson diversity index B) Fungi Shannon diversity index. C) Plant Simpson diversity index D) Plant Shannon diversity index. Letters above the bars indicate significant groupings as determined by Tukey's HSD post hoc test and error bars show standard error of the mean. NS indicates no significant treatment effect. Note that Shannon and Simpson indices are scaled inversely (i.e. higher index = lower diversity).

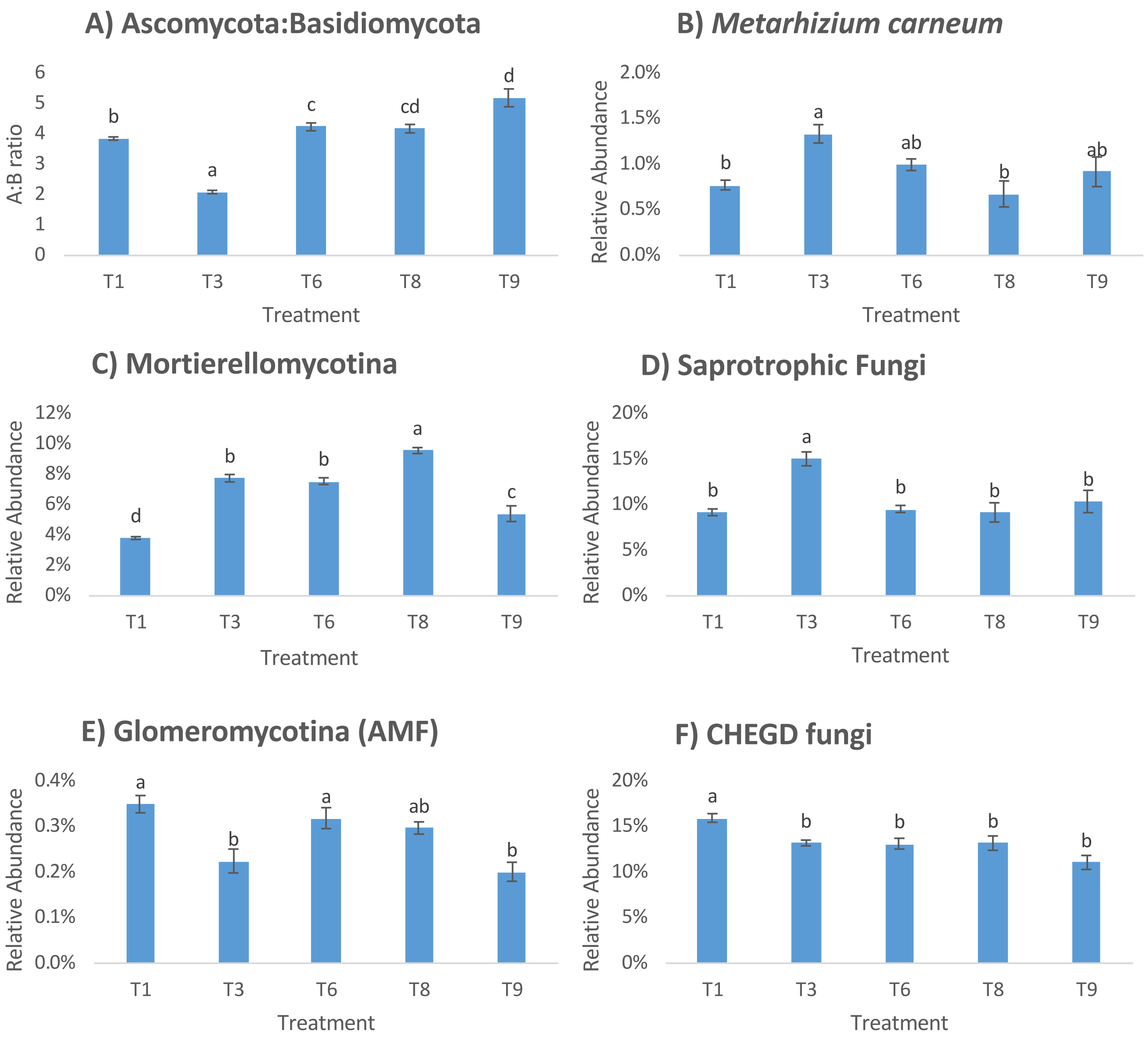


**SuppData 7.** Relative abundance of fungal groups by storage treatment for the alluvial soil (Gogerddan) A) Ascomycota to Basidiomycota ratio; B) Metarhizium carneum; C) Mortierellomycota; D) Saprotrophic fungi; E) Glomeromycotina (Arbuscular mycorrhizal fungi); F) Grassland fungi (CHEGD). Letters above the bars indicate significant groupings as determined by Tukey's HSD post hoc test and error bars show standard error of the mean.
